# *Saccharomyces* Genome Database: Advances in Genome Annotation, Expanded Biochemical Pathways, and Other Key Enhancements

**DOI:** 10.1101/2024.09.16.613348

**Authors:** Stacia R. Engel, Suzi Aleksander, Robert S. Nash, Edith D. Wong, Shuai Weng, Stuart R. Miyasato, Gavin Sherlock, J. Michael Cherry

## Abstract

Budding yeast (*Saccharomyces cerevisiae*) is the most extensively characterized eukaryotic model organism and has long been used to gain insight into the fundamentals of genetics, cellular biology, and the functions of specific genes and proteins. The *Saccharomyces* Genome Database (SGD) is a scientific resource that provides information about the genome and biology of *S. cerevisiae*. For more than 30 years, SGD has maintained the genetic nomenclature, chromosome maps, and functional annotation for budding yeast along with search and analysis tools to explore these data. Here we describe recent updates at SGD, including the two most recent reference genome annotation updates, expanded biochemical pathways representation, changes to SGD search and data files, and other enhancements to the SGD website and user interface. These activities are part of our continuing effort to promote insights gained from yeast to enable the discovery of functional relationships between sequence and gene products in fungi and higher eukaryotes.

## INTRODUCTION

The *Saccharomyces* Genome Database (SGD; https://www.yeastgenome.org) is a scientific knowledgebase that provides comprehensive and up-to-date information about the genome and biology of the yeast *Saccharomyces cerevisiae*. It serves as a valuable resource for researchers studying yeast biology and genetics by offering information on genes, proteins, pathways, and phenotypes. Scientists can use SGD to explore the functions of genes, track genetic and physical interactions, and access curated literature related to yeast genetics and genomics. SGD plays a crucial role in advancing our understanding of molecular mechanisms and processes in yeast and serves as a central repository for yeast-related data and information.

Since 1993, SGD has been assembling and cataloging scientific data regarding the genome and proteome of budding yeast, and distributing that information to the public via an open-access web interface and download service. Budding yeast data that can be found at SGD include the reference genome (Engel et al. 2022), which is a single consensus representative *S. cerevisiae* genome sequence against which all other sequences can be compared, and various analysis tools (Balakrishnan et al. 2004, Cherry 2015, Christie et al. 2004, Hirschman et al. 2006, Sheppard et al. 2016) and data files that allow interrogation of the genome sequence and its products for a wide variety of applications (Engel et al. 2018, Hellerstedt et al. 2017, Ng et al. 2019, Wong et al. 2019).

Yeast have proven especially useful for studying various aspects of biology, including the regulation of gene expression through upstream open reading frames (uORFs; Blank et al. 2017, Cartwright et al. 2017, Vindu et al. 2021, Yang et al. 2023), emergence of newly evolved genes (Chang et al. 2023, Wacholder and Carvunis 2023, Wacholder et al. 2023), complex gene structure (Balarezo-Cisneros et al. 2021, Feng et al. 2022, Xu et al. 2009, Yang et al. 2023), and cellular metabolism (Ljungdahl and Daignan-Fornier 2012, Thomas and Surdin-Kerjan 1997). The conservation of many molecular mechanisms between yeast and higher eukaryotes makes findings from yeast studies broadly applicable to understanding the biology of more complex organisms, including humans. Studying these processes in yeast can help uncover basic biological principles that are widely applicable, both to general understanding and for solutions to specific questions. Findings from yeast studies can be validated in other model organisms or human cell lines to assess conservation and relevance across species.

With its simple and well-characterized genome, *S. cerevisiae* makes it easier to study gene functions, regulatory elements, and cellular processes. Advances in yeast research, such as the development of molecular biology techniques, genetic tools, and resources like SGD make yeast a valuable model organism for transferring knowledge to other species by providing a simpler and more tractable system to study fundamental biological processes, test hypotheses, and develop experimental techniques. Here we describe recent changes at SGD, including the two most recent reference genome annotation updates, expanded biochemical pathways representation, improvements to SGD search and data files, and other enhancements to the SGD website and user interface.

## GENOME ANNOTATION UPDATES

### Genome version R64.4.1

In the first of two recent updates, the *S. cerevisiae* strain S288C reference genome annotation was updated to release R64.4.1, dated 2023-08-23 (Table 1). The underlying genome sequence itself was not altered in any way. The update included the addition of eight noncoding RNAs (ncRNAs) and the addition of three new upstream ORFs (uORFs). Three open reading frames (ORFs) were demoted from ‘Uncharacterized’ to ‘Dubious’ because they were found to overlap tRNAs, have multiple frameshifts and/or indels in the coding region, and had minimal evidence to support their existence.

**Table 1.** The *S. cerevisiae* strain S288C reference genome annotation was updated to release R64.4.1, dated 2023-08-23.

| Chromosome | Feature | Description of change | Reference |
| --- | --- | --- | --- |
| III | SUT035 / YNCC0015W | New ncRNA<br>chrIII:205766..205942 | Xu et al. 2009, Balarezo-Cisneros et al. 2021 |
| IV | YDR278C | Change ORF qualifier from Uncharacterized to Dubious | Requested by NCBI |
| IV | SUT053 / YNCD0033W | New ncRNA<br>chrIV:506334..507774 | Xu et al. 2009, Balarezo-Cisneros et al. 2021 |
| IV | SUT468 / YNCD0034C | New ncRNA<br>chrIV:506546..507450 | Xu et al. 2009, Balarezo-Cisneros et al. 2021 |
| VII | SUT532 / YNCG0047C | New ncRNA<br>chrVII:17213..17709 | Xu et al. 2009, Balarezo-Cisneros et al. 2021 |
| VII | SUT125 / YNCG0048W | New ncRNA<br>chrVII:650855..651159 | Xu et al. 2009, Balarezo-Cisneros et al. 2021, Feng et al. 2022 |
| VII | SUT126 / YNCG0049W | New ncRNA<br>chrVII:660087..661399 | Xu et al. 2009, Balarezo-Cisneros et al. 2021 |
| XII | *FPS1* /<br>*YLL043W* | New uORF<br>chrXII:49924..49932 | Cartwright et al. 2017 |
| XIV | *ACC1* /<br>*YNR016C* | New uORF<br>chrXIV:661704..661715 | Blank et al. 2017 |
| XIV | *HOL1* /<br>*YNR055C* | New uORF<br>chrXIV:730381..730401 | Vindu et al. 2021 |
| XV | *YOL013W-A* | Change ORF qualifier from<br>Uncharacterized to Dubious | Requested by NCBI |
| XVI | *SUT390* /<br>*YNCP0025W* | New ncRNA<br>chrXVI:52977..53465 | Xu et al. 2009, Feng et al. 2022 |
| XVI | *SUT418* /<br>*YNCP0026W* | New ncRNA<br>chrXVI:588998..589830 | Xu et al. 2009, Feng et al. 2022 |
| XVI | *YPR108W-A* | Change ORF qualifier from<br>Uncharacterized to Dubious | Requested by NCBI |

### Genome version R64.5.1

In the second of two recent updates, the *S. cerevisiae* strain S288C reference genome annotation was updated to release R64.5.1, dated 2024-05-29 (Table 2). Once again, the underlying genome sequence was not altered; the chromosome sequences remain stable and unchanged. The update included the addition of six new ORFs and six new uORFs, a shifted start for one ORF, and the upgrade of one ORF from Dubious to Verified because a stable translation product was detected.

**Table 2.** The *S. cerevisiae* strain S288C reference genome annotation was updated to release R64.5.1, dated 2024-05-29.

| Chromosome | Feature | Description of change | Reference |
| --- | --- | --- | --- |
| II | *ATG12* /<br>*YBR217W* | New uORF chrII:657824..657835,<br>partially overlaps CDS | Yang et al. 2023 |
| IV | *YDL204W-A* | New ORF chrIV:94133..94285 | Wacholder et al. 2023 |
| VI | *YFR035W-A* | New ORF chrVI:226260..226550 | Wacholder and Carvunis 2023 |
| VII | *YGR016C-A* | New ORF chrVII:523353..523246 | Wacholder et al. 2023, Chang et al. 2023 |
| IX | *EFM4* /<br>*YIL064W* | Move start 84 nucleotides<br>downstream, new coordinates<br>chrIX:242027..242716 | Hamey et al. 2024 |
| IX | *YIL059C* | Change ORF qualifier from<br>Dubious to Verified | Wacholder and Carvunis 2023 |
| XIII | *YMR106W-A* | New ORF chrXIII:480924..481187 | Wacholder and Carvunis 2023 |
| XIV | *YNL040C-A* | New ORF chrXIV:552558..552478 | Wacholder et al. 2023 |
| XIV | *YNL155C-A* | New ORF chrXIV:342135..341911 | Wacholder and Carvunis 2023 |
| XV | ATG19 / YOL082W | New uORF chrXV:168632..168679 | Yang et al. 2023 |
| XVI | ATG5 / YPL149W | 4 new uORFs:<br>chrXVI:271236..271277,<br>chrXVI:271252..271302,<br>chrXVI:271299..271307,<br>chrXVI:271302..271307 | Yang et al. 2023 |
| XVI | ATG13 / YPR185W | New uORF<br>chrXVI:907211..907351, partially<br>overlaps CDS | Yang et al. 2023 |

## CHANGES TO *SACCHAROMYCES CEREVISIAE* GFF FILE

The saccharomyces_cerevisiae.gff contains data regarding sequence features of *S. cerevisiae* strain S288C and related information such as locus descriptions and Gene Ontology (The Gene Ontology Consortium 2023) annotations. It is fully compliant with Generic Feature Format Version 3 (https://gmod.org/wiki/GFF3.html), and is updated weekly. This is a standard format used by many genomics and database groups. SGD uses the GFF file to load the reference data tracks into SGD’s genome browser resource (https://jbrowse.yeastgenome.org).

After November 2020, SGD updated the transcript features in the GFF file to reflect experimentally determined transcripts (Pelechano et al. 2013, Ng et al. 2020), when possible. The longest transcripts were determined for two different widely-used growth media - galactose and dextrose. When available, experimentally determined transcripts for one or both conditions were added for a gene. Where these data were absent, transcript entries matching the start and stop coordinates of the ORF were used.

In February 2024, SGD edited the ‘gene’ entries in the file to extend the coordinates to encompass the start and stop coordinates of the longest experimentally determined transcripts, regardless of condition. This change was made in order to comply with JBrowse 2 (Diesh et al. 2023), a newer and more extensible genome browser, which requires that ‘gene’ features in GFF files represent a longer region than the features that make up a ‘gene’ (coding sequences, mRNA, etc.).

## BIOCHEMICAL PATHWAYS

SGD’s YeastPathways (https://pathway.yeastgenome.org; Cherry 2015) is a database of 220 conserved metabolic pathways and their corresponding enzymes in *S. cerevisiae*, manually curated and maintained by the curation team at SGD. YeastPathways enables visualization of yeast metabolism from large metabolic networks to individual pathways, and from biochemical reactions down to individual metabolites. Search tools and click-to-browse features in YeastPathways enable quick navigation and intuitive exploration of yeast metabolism.

We recently completed a major update to the YeastPathways content. As the first major update since 2012, we updated 62 pathways with expert summaries on pathway genetics, biochemistry, regulation, and more. Thirty-three new pathways with specificity for yeast biochemistry were propagated from MetaCyc at SRI (Caspi R, et al. 2018), and 105 existing pathways were edited for proper enzymatic classification, reaction connectivity, and gene attribution. Compounds that were previously missing a chemical structure have also now been updated, along with the stoichiometry and scheme of many pathway reactions.

Because many fundamental molecular processes and pathways are evolutionarily conserved between yeast and higher eukaryotes, including humans, the curated metabolic pathway information has great value for the transfer of knowledge to other organisms. Therefore, the YeastPathways data were exported in BioPAX (Demir et al. 2010) format for import into Noctua, a tool for collaborative curation of biological pathways and gene annotations that was developed by the GO Consortium (Thomas et al. 2019). BioPAX provides a standardized format for representing biological pathways, allowing researchers to integrate pathway information from different sources and databases. Noctua can import pathway data encoded in BioPAX format to populate the pathway editor with molecular interactions, biological processes, and regulatory relationships, and can utilize BioPAX files to combine pathway data from multiple datasets for pathway curation and analysis. Pathways curated and edited in Noctua can be exported both as GO annotations for yeast and orthologous genes in other species, or as pathway annotations in BioPAX, which facilitates sharing of curated pathways with other researchers, databases, and pathway analysis tools using a standard format, promoting data exchange and collaboration within the scientific community.

**Figure 1.**
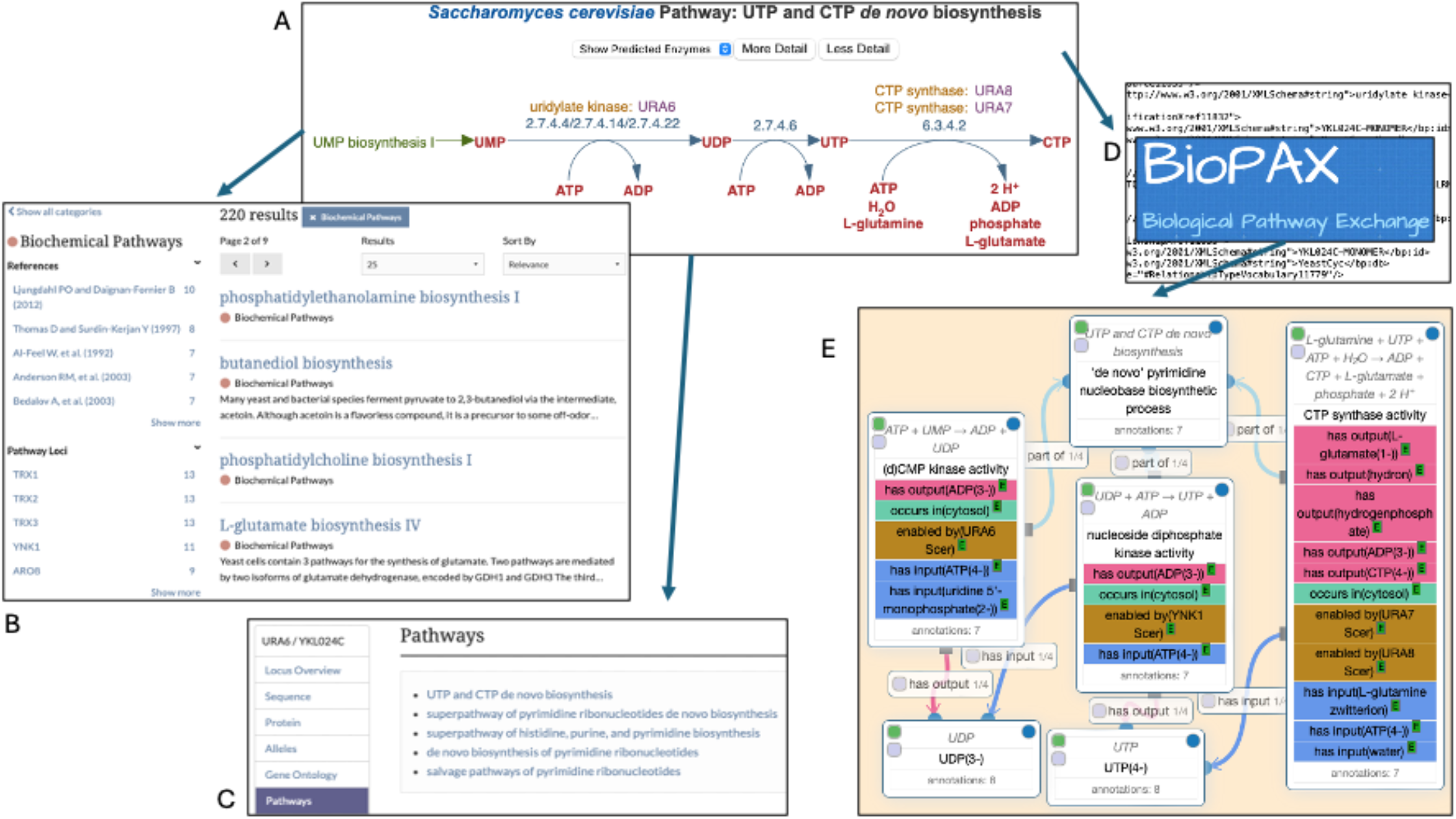
Curated metabolic pathways from YeastPathways (A; UTP and CTP *de novo* biosynthesis, https://pathway.yeastgenome.org/YEAST/NEW-IMAGE?type=PATHWAY&object=PWY-7176) are accessible via SGD Search (B; https://www.yeastgenome.org/search) and SGD Gene pages (C; *URA6*, https://www.yeastgenome.org/locus/URA6). Data from YeastPathways have been exported via BioPAX format (D) to create Gene Ontology annotations using the Noctua collaborative curation tool for pathways and gene annotations (E; UTP and CTP *de novo* biosynthesis, http://noctua.geneontology.org/editor/graph/gomodel:YeastPathways_PWY-7176).

YeastPathways can be accessed via the Function menu in the purple toolbar that runs across the top of most SGD webpages or from the Pathways section on SGD Gene pages. To make the pathways more readily accessible, we added the pathways to SGD Search. The category “Biochemical Pathways” is now available, with facets (i.e., subcategories) for References and Loci. For even easier access, we also added the Pathway names and IDs to the autocomplete in the Search box, to enable quick browsing.

## UPDATES TO SGD SEARCH

Because utilizing the SGD search box provides the most efficient and direct access to the content on the site, we have recently added new data and modified existing data mappings to optimize search performance and capabilities. We have added a new category for Datasets, with over 3,700 yeast datasets accessible for searching by reference, keyword, assay, and lab. A new Strains subcategory has been added to the Reference search. Macromolecular complexes can now be searched with aliases, reference, subunit, function, process, and location. Alleles can be searched via their descriptions, SGDIDs, reference, allele type, gene, and phenotype. RNA products can now be searched using RNAcentral IDs. The improved search functionality enhances the user experience and increases user satisfaction through improved navigation which provides easier access to information, higher relevance in search results, improved data retrieval, and overall better efficiency.

## OTHER UPDATES TO THE WEB INTERFACE

We regularly update the SGD web interface to enhance user experience, improve visual appeal, incorporate new features, align with modern design trends, increase usability, and improve user engagement. The modifications and enhancements described below make the website more user-friendly and effective without the implementation of major overhauls or revisions.

SGD biocurators use the Chemical Entities of Biological Interest (ChEBI) Ontology (Hastings et al. 2016), maintained by EMBL-EBI, to describe chemicals used in experiments curated from yeast publications and displayed on SGD webpages. We recently added chemical structures provided by ChEBI to the Chemical pages in SGD.

In 2011, SGD implemented InterMine (http://www.InterMine.org; Smith et al. 2012), an open source data warehouse system with a sophisticated querying interface, to create YeastMine (Balakrishnan et al. 2012, Hellerstedt et al. 2017), a multifaceted search and retrieval environment that provided access to diverse data types. YeastMine served as a powerful search interface, a discovery tool, a curation aid, and a complex database presentation format. We recently moved the YeastMine data into AllianceMine, hosted by the Alliance of Genome Resources (Alliance of Genome Resources Consortium 2024), of which SGD is a founding member. Users can get started with AllianceMine by going to the Templates page, and filtering by the category ‘YeastMine’. The data from YeastMine are also available on the SGD Downloads site (http://sgd-archive.yeastgenome.org). Information regarding genes and IDs, etc., are in the chromosomal_features directory, and a variety of annotation files for different types of data can be found in the literature directory.

The implementation of Textpresso (Müller et al. 2004) by SGD has recently been updated. Each week, SGD biocurators triage new publications from PubMed to load the newest yeast papers into the database. Once they have been added into SGD, those papers get indexed and loaded into Textpresso, a tool for full-text mining and searching, which provides results shown in the context of the full text, with matches to query terms highlighted *in situ*. Textpresso allows several user-friendly options, including use of Boolean operators, custom corpus creation allowing users to decide which papers to search, search scope options for document or sentence, and search location options for constraining searches to specific sections of papers. Content updates in SGD’s Textpresso are now happening on a weekly basis, enabling full-text search of the very latest yeast papers added to SGD. Textpresso can be accessed via the “Full-text Search” link under “Literature” in the purple toolbar that runs across the top of most SGD webpages.

SGD was recently chosen as a Global Core Biodata Resource (GCBR; https://globalbiodata.org/what-we-do/global-core-biodata-resources) in recognition of our commitment to providing high-quality and valuable biological data to the global research community. We are honored to be selected as a GCBR, and we are dedicated to upholding the highest standards of data integrity, accessibility, and usability to support cutting-edge research and scientific discovery on a global scale. This recognition motivates us to continue expanding and improving SGD to empower researchers worldwide in advancing knowledge and innovation in yeast genetics, genomics, and the life sciences as a whole.

## FUTURE DIRECTIONS

SGD plays a crucial role in organizing, curating, and disseminating biological information related to the model organism budding yeast *S. cerevisiae*. Because many fundamental molecular processes and pathways are evolutionarily conserved between yeast and higher eukaryotes, *S. cerevisiae* is highly useful for transferring that knowledge to other organisms. As one of seven founders of the Alliance of Genome Resources (Alliance of Genome References Consortium 2024), a new central knowledgebase for *Saccharomyces cerevisiae* (yeast), *Caenorhabditis elegans* (worm), *Drosophila melanogaster* (fly), *Danio rerio* (zebrafish), *Xenopus laevis* (frog), *Rattus norvegicus* (rat), *Mus musculus* (mouse), and *Homo sapiens* (human), SGD is positioned to continue advancing scientific research and supporting the needs of the scientific community. Adopting and promoting data standards and interoperable formats will facilitate data exchange and integration between different model organism databases and biological resources. Ensuring data consistency and compatibility enables seamless collaboration and cross-referencing of information across research communities.

As such, we will continue our work with this consortium to harmonize common data types and create a unified web resource. Integrating data from various sources allows researchers to explore complex biological relationships and gain comprehensive insights into gene function and regulation. A large amount of this work has been completed, and integration proceeds apace. SGD’s JBrowse genome browser, YeastMine data warehouse, and Textpresso full-text search tool have already been incorporated into the Alliance of Genome Resources. Current efforts include an integrated BLAST tool based on SequenceServer (https://sequenceserver.com), which we hope to release later this year.

## DATA AVAILABILITY

All SGD data and tools are freely available at https://www.yeastgenome.org. The SGD API is freely available at https://www.yeastgenome.org/api/doc. SGD downloads are freely available at http://sgd-archive.yeastgenome.org. YeastMine data within AllianceMine are freely available at https://www.alliancegenome.org/alliancemine.

## ACKNOWLEDGEMENTS

SGD thanks the yeast community for their invaluable contribution of data and for providing valuable feedback on the SGD website and community needs. We thank the SGD Advisory Board for their continued support. We thank Benjamin Good and Dustin Ebert for their assistance with importing the YeastPathways data into Noctua.

## REFERENCES

Alliance of Genome Resources Consortium. 2024. Updates to the Alliance of Genome Resources central infrastructure. Genetics. 227(1):iyae049.

Balakrishnan R, Christie KR, Costanzo MC, Dolinski K, Dwight SS, Engel SR, Fisk DG, Hirschman JE, Hong EL, Nash R, et al. 2005. Fungal BLAST and model organism BLASTP best hits: new comparison resources at the Saccharomyces Genome Database (SGD). Nucleic Acids Res. 33:D374–D377.

Balakrishnan R, Park J, Karra K, Hitz BC, Binkley G, Hong EL, Sullivan J, Micklem G, Cherry JM (2012) YeastMine - An integrated data warehouse for S. cerevisiae data as a multi-purpose tool-kit. Database (Oxford). Mar 20:2012:bar062.

Balarezo-Cisneros LN, Parker S, Fraczek MG, Timouma S, Wang P, O’Keefe RT, Millar CB, Delneri D. 2021. Functional and transcriptional profiling of non-coding RNAs in yeast reveal context-dependent phenotypes and in trans effects on the protein regulatory network. PLoS Genet. 17(1):e1008761.

Blank HM, Perez R, He C, Maitra N, Metz R, Hill J, Lin Y, Johnson CD, Bankaitis VA, Kennedy BK, et al. 2017. Translational control of lipogenic enzymes in the cell cycle of synchronous, growing yeast cells. EMBO J. 36(4):487–502.

Cartwright SP, Darby RAJ, Sarkar D, Bonander N, Gross SR, Ashe MP, Bill RM. 2017. Constitutively-stressed yeast strains are high-yielding for recombinant Fps1: implications for the translational regulation of an aquaporin. Microb Cell Fact. 16(1):41.

Caspi R, Billington R, Fulcher CA, Keseler IM, Kothari A, Krummenacker M, Latendresse M, Midford PE, Ong Q, Ong WK, et al. 2018. The MetaCyc database of metabolic pathways and enzymes. Nucleic Acids Res. 46(D1):633–D639.

Chang S, Joyson M, Kelly A, Tang L, Iannotta J, Rich A, Coelho NC, Carvunia AR. 2023. Unannotated Open Reading Frame in Saccharomyces cerevisiae Encodes Protein Localizing to the Endoplasmic Reticulum. MicroPubl Biol. Oct 20:2023:10.17912/micropub.biology.000992.

Cherry JM. 2015. The Saccharomyces Genome Database: Exploring biochemical pathways and mutant phenotypes. Cold Spring Harb Protoc. 2015(12):pdb.prot088898.

Christie KR, Weng S, Balakrishnan R, Costanzo MC, Dolinski K, Dwight SS, Engel SR, Feierbach B, Fisk DG, Hirschman JE, et al. 2004. Saccharomyces Genome Database (SGD) provides tools to identify and analyze sequences from Saccharomyces cerevisiae and related sequences from other organisms. Nucleic Acids Res. 32:D311–D314.

Demir E, Cary MP, Paley S, Lemur C, Vastrik I, Wu G, D’Eustachio P, Schaefer C, Luciano J, Schacherer F, et al. 2010. The BioPAX community standard for pathway data sharing. Nat Biotechnol. 28(9):935–42.

Diesh C, Stevens GJ, Xie P, De Jesus Martinez T, Hershberg EA, Leung A, Guo E, Dider S, Zhang J, Bridge C, et al. 2023. JBrowse 2: a modular genome browser with views of synteny and structural variation. Genome Biol. 17;24(1):74.

Engel SR, Skrzypek MS, Hellerstedt ST, Wong ED, Nash RS, Weng S, Binkley Sheppart TK, Karra K, Cherry JM. 2018. Updated regulation curation model at the Saccharomyces Genome Database. Database (Oxford). 2018:bay007. doi: 10.1093/database/bay007.

Engel SR, Wong ED, Nash RS, Aleksander S, Alexander M, Douglass E, Karra K, Miyasato SR, Simison M, Skrzypek MS, et al. 2022. New data and collaborations at the Saccharomyces Genome Database: updated reference genome, alleles, and the Alliance of Genome Resources. Genetics. 220(4):iyab224.

Feng MW, Delneri D, Millar CB, O’Keefe RT. 2022. Eisosome disruption by noncoding RNA deletion increases protein secretion in yeast. PNAS Nexus 1(5):pgac241.

The Gene Ontology Consortium. 2023. The Gene Ontology knowledgebase in 2023. Genetics. 224(1):iyad031.

Hamey JJ, Nguyen A, Haddad M, Vázquez-Campos X, Pfeiffer PG, Wilkins MR. 2024. Methylation of elongation factor 1A by yeast Efm4 or human eEF1A-KMT2 involves a beta-hairpin recognition motif and crosstalks with phosphorylation. J Biol Chem. 300(2):105639.

Hastings J, Owen G, Dekker A, Ennis M, Kale N, Muthukrishnan V, Turner S, Swainston N, Mendes P, Steinbeck C. 2016. ChEBI in 2016: Improved services and an expanding collection of metabolites. Nucleic Acids Res.

Hellerstedt ST, Nash RS, Weng S, Paskov KM, Wong ED, Karra K, Engel SR, Cherry JM. 2017. Curated protein information in the Saccharomyces Genome Database. 2017:bax011. doi: 10.1093/database/bax011.

Hirschman JE, Balakrishnan R, Christie KR, Costanzo MC, Dwight SS, Engel SR, Fisk DG, Hong EL, Livstone MS, Nash R, et al. 2006. Genome snapshot: a new resource at the Saccharomyces Genome Database (SGD) presenting an overview of the Saccharomyces cerevisiae genome. Nucleic Acids Res. 34:D442–D445.

Ljungdahl PO, Daignan-Fornier B. 2012. Regulation of amino acid, nucleotide, and phosphate metabolism in Saccharomyces cerevisiae. Genetics. 190(3):885–929.

Müller HM, Kenny EE, Sternberg PW. 2004. Textpresso: an ontology-based information retrieval and extraction system for biological literature. PLoS Biol. 2(11):e309.

Ng PC, Wong ED, MacPherson KA, Aleksander S, Argasinska J, Dunn B, Nash RS, Skrzypek MS, Gondwe F, Jha S, et al. 2020. Transcriptome visualization and data availability at the Saccharomyces Genome Database. Nucleic Acids Res. 48(D1):D743–D748.

Pelechano V, Wei W, Steinmetz LM. 2013. Extensive transcriptional heterogeneity revealed by isoform profiling. Nature. 497:127–131.

Smith RN, Aleksic J, Butano D, Carr A, Contrino S, Hu F, Lyne M, Lyne R, Kalderimis A, Rutherford K, et al. 2012. InterMine: a flexible data warehouse system for the integration and analysis of heterogeneous biological data. Bioinformatics. 28(23):3163–5.

Sheppard TK, Hitz BC, Engel SR, Song G, Balakrishnan R, Binkley G, Costanzo MC, Dalusag KS, Demeter J, Hellerstedt S, et al. 2016. The Saccharomyces Genome Database Variant Viewer. Nucleic Acids Res.44:D698–D702.

Thomas D, Surdin-Kerjan Y. 1997. Metabolism of sulfur amino acids in Saccharomyces cerevisiae. Microbiol Mol Biol Rev. 61(4):503–32.

Thomas PD, Hill DP, Mi H, Osumi-Sutherland D, Van Auken K, Carbon S, Balhoff JP, Albou LP, Good B, Gaudet P, et al. 2019. Gene Ontology Causal Activity Modeling (GO-CAM) moves beyond GO annotations to structured descriptions of biological functions and systems. Nat Genet. 51(10):1429–1433.

Vindu A, Shin BS, Choi K, Christenson ET, Ivanov IP, Cao C, Banerjee A, Dever TE. 2021. Translational autoregulation of the S. cerevisiae high-affinity polyamine transporter Hol1. Mol Cell. 81(19):3904-3918.e6.

Wacholder A, Carvunis AR. 2023. Biological factors and statistical limitations prevent detection of most noncanonical proteins by mass spectrometry. PLoS Biol. 21(12):e3002409.

Wacholder A, Parikh SB, Coelho NC, Acar O, Houghton C, Chou L, Carvunis AR. 2023. A vast evolutionarily transient translatome contributes to phenotype and fitness. Cell Syst. 14(5):363-381.e8.

Wong ED, Skrzypek MS, Weng S, Binkley G, Meldal BHM, Perfetto L, Orchard SE, Engel SR, Cherry JM. 2019. Integration of macromolecular complex data into the Saccharomyces Genome Database. Database. 2019:baz008.

Xu Z, Wei W, Gagneur J, Perocchi F, Clauder-Münster S, Camblong J, Guffanti E, Stutz F, Huber W, Steinmetz LM. 2009. Bidirectional promoters generate pervasive transcription in yeast. Nature 457(7232):1033–7.

Yang Y, Gatica D, Liu X, Wu R, Kang R, Tang D, Klionsky DJ. 2023. Upstream open reading frames mediate autophagy-related protein translation. Autophagy 19(2):457–473.

